# Persistent fatty acid catabolism during plant oil synthesis

**DOI:** 10.1101/2025.02.03.636323

**Authors:** Somnath Koley, Poonam Jyoti, Maneesh Lingwan, Michael Wei, Chunhui Xu, Kevin L. Chu, Russell B. Williams, Abraham J. Koo, Jay J. Thelen, Dong Xu, Doug K. Allen

## Abstract

Plant lipids are an essential energy source for diets and are a sustainable alternative to petroleum-based fuels and feed stocks. Fatty acid breakdown during seed germination is crucial for seedling establishment but unexpected during seed filling. Here, we demonstrate that the simultaneous biosynthesis and degradation of fatty acids begins early and continues across all phases of oil filling and throughout the photoperiod. Tests in camelina, rapeseed and engineered high-oil tobacco line confirmed that concomitant synthesis and breakdown in oil-producing tissues over development is the rule rather than the exception. We show that transgenics, designed for significantly higher fatty acid biosynthesis, fail to achieve a proportional increase in storage lipid levels due to increasing degradation, potentially explaining the underperformance of engineered lines compared to expectations.

## INTRODUCTION

Lipids are a highly desirable renewable feed stock for fuels and renewable feed stocks as part of a sustainable bioeconomy because they are energetically dense, containing twice the energy of carbohydrates per gram ^1–3^. The demand for lipids is expected to double by 2050; however, renewable sources such as oilseeds are currently insufficient to fulfill the needs. Efforts to increase oil levels in seeds have focused on enhancing fatty acid ^4^ and lipid biosynthesis ^5–7^, or altering the supply of carbon precursors ^8^, but have yet to result in anticipated transformative gains, suggesting that other processes contribute to final oil concentrations. One possibility is that engineered increases in lipid production are offset by enhanced fatty acid breakdown. Without measurements that can distinguish synthesis and turnover processes, fatty acid degradation in oil-filling tissues could go easily undetected due to the high rates of fatty acid biosynthesis, resulting in billions of dollars in lost potential productivity worldwide ^9^ (Text S1).

The mobilization of stored energy in lipids is also crucial to sustain living systems. Animals rely on lipid turnover to survive cold winters or periods of hibernation. Without the breakdown of fatty acids, humans would become obese, and seeds would fail to germinate, leading to barren tracts of land and the collapse of life on Earth. Germination in plants involves the breakdown of lipid-derived fatty acids in the peroxisome to generate carbon skeletons and energy that sustain the growth of seedlings. Apart from germination, significant oxidation of fatty acids is thought to be limited to maturation of seeds and leaves, including those that have been genetically altered ^10–14^, although the breakdown process may be active at other times in plant tissues to detoxify cells ^15–17^. Lipid remodeling and breakdown to adjust membrane lipid composition, lipid signaling or jasmonic acid synthesis has been extensively studied under stress ^18–20^, but the quantitative breakdown of fatty acids originating from lipids that may occur as a part of regular growth ^21^ when significant oil is accumulated has received less attention. Assessing the concomitant biosynthesis and breakdown of fatty acids would not be possible without isotopes ^22,23^.

The breakdown of fatty acids proceeds by biochemical steps that are essentially the reverse of two-carbon additions during fatty acid biosynthesis. Instead of acetyl groups added on an acyl-carrier protein (ACP) backbone in the chloroplast during biosynthesis ^24,25^, they are released from an acyl-chain attached to coenzyme A (CoA) during fatty acid breakdown, known as β-oxidation ^21^ (Figure 1A). The process was first observed in dogs ^26^ and detailed later in plant glyoxosomes or peroxisomes in association with glyoxylate activity in seedlings ^27^. Long-chain acyl-CoAs (≥C16 in acyl-chain) can result from biosynthetic or breakdown activities. Shorter-chain (C4-14) acyl-CoAs are specific to β-oxidation, with notable exceptions that include plants that produce lipids containing medium-chain fatty acids (e.g., *Cocos nucifera*, *Cuphea sp.* and *Umbellularia californica*) ^28^. Acyl-CoA oxidase (ACX1-4) activities involved in the breakdown of specific chain lengths have been documented ^15,29–32^ as have the specific enzymes involved in handling double bonds originating from polyunsaturated fatty acids ^20,21,33,34^. The intermediates of β-oxidation include enoyl and ketoacyl-CoAs that are also associated with cytosolic fatty acid elongation for chains of 20 carbons or longer; however, shorter length intermediates are exclusively involved in β-oxidation. Further, unsaturated shorter-chain CoAs are specifically derived from polyunsaturated fatty acids (Figure 1B) and unequivocally indicate lipid degradation because the process of adding multiple double bonds to form polyunsaturated fatty acids occurs only on a lipid backbone in plants ^12,35,36^ (Figure 1A) but have not been quantified.

**Figure 1.**
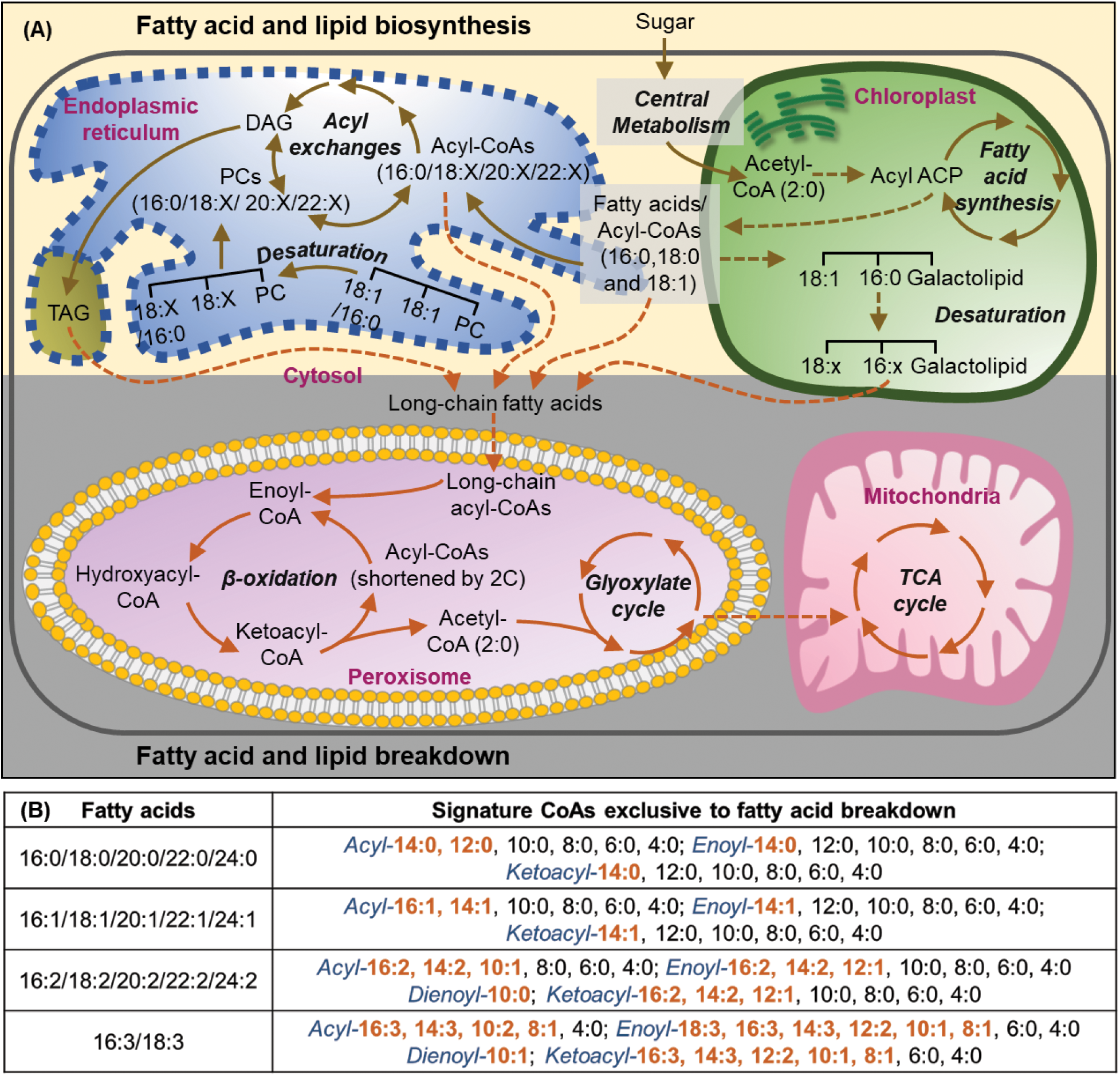
Fatty acid and lipid metabolism in plants (A) Fatty acid and lipid biosynthesis occur in the chloroplast and endoplasmic reticulum (ER), respectively. Generation of polyunsaturated fatty acids takes place on a lipid backbone. Fatty acid breakdown occurs in the peroxisomes of plants. Acetyl-CoA, which is produced during fatty acid oxidation, is converted into organic acids via the glyoxylate cycle before being transported to the mitochondria for further metabolism. (B) Short- to medium-chain CoAs are characteristic of breakdown of different fatty acids. Breakdown-specific CoAs in orange color are specific to fatty acid substrates. Abbreviations (also listed in Data S1): TAG, triacylglycerol; DAG, diacylglycerol; ACP, acyl-carrier protein; PC, phosphatidylcholine; x, polyunsaturation.

Saturated acyl-CoAs associated with β-oxidation in germinating seedlings were measured ^30,31,37^, but the levels of oxidation in other plant tissues including developing seeds that are concentrated in oil and provide the economic value of crops remain unclear. Here, we present multiple lines of evidence that fatty acid oxidation occurs concomitantly with oil synthesis in oil-rich tissues at all oil-filling stages. Synthesis and breakdown of fatty acids and lipids are precisely controlled throughout the photoperiod. Additionally, β-oxidation occurs at varied levels in engineered lines that were modified to increase oil.

## RESULTS AND DISCUSSION

We developed a mass spectrometry method to sensitively measure acyl-CoAs extending descriptions in literature ^37^ (Text S2) to analyze intermediates in the oxidation cycle originating from breakdown of saturated and unsaturated fatty acids (Fig. 1B). The method was initially tested in germinating camelina seedlings since fatty acid breakdown is an essential process of germination in oilseeds. The presence of shorter-chain unsaturated acyl-CoAs, which have not been previously measured in plants, indicated conclusively that lipid breakdown occurred. Saturated and unsaturated forms of acyl-, enoyl-, dienoyl-, ketoacyl-CoAs were detected that are signatures of fatty acid oxidation (Text S2, Figure S1).

### Fatty acid biosynthesis and breakdown is a synchronized process

As seeds are the singular plant propagule with significant concentrations of lipids, even a small amount of fatty acid recycling during seed filling could dramatically affect the final composition; however, the possibility of this perceived futile cycle remains an open question. To assess the breakdown process when significant fatty acid biosynthesis occurs, three developmental stages were measured in camelina (15, 20 and 25 days post anthesis or DPA) and rapeseed (15, 25 and 35 DPA) during seed development and also in leaves of transgenic high-oil tobacco ^38^ (30, 45 and 60 days after sowing or DAS). The stages were defined to capture oil biosynthesis after initiation (the 1^st^ stage; early), at the highest rate of oil biosynthesis (the 2^nd^ stage; mid), and at the final phase of filling (the 3^rd^ stage; late) when tissues were still green and prior to desiccation or senescence (Figure S2). Fifty-one esterified CoAs were quantified including saturated and unsaturated forms of acyl-, enoyl- and ketoacyl-CoAs, of which 34 were specific to breakdown processes (Figure 2).

**Figure 2.**
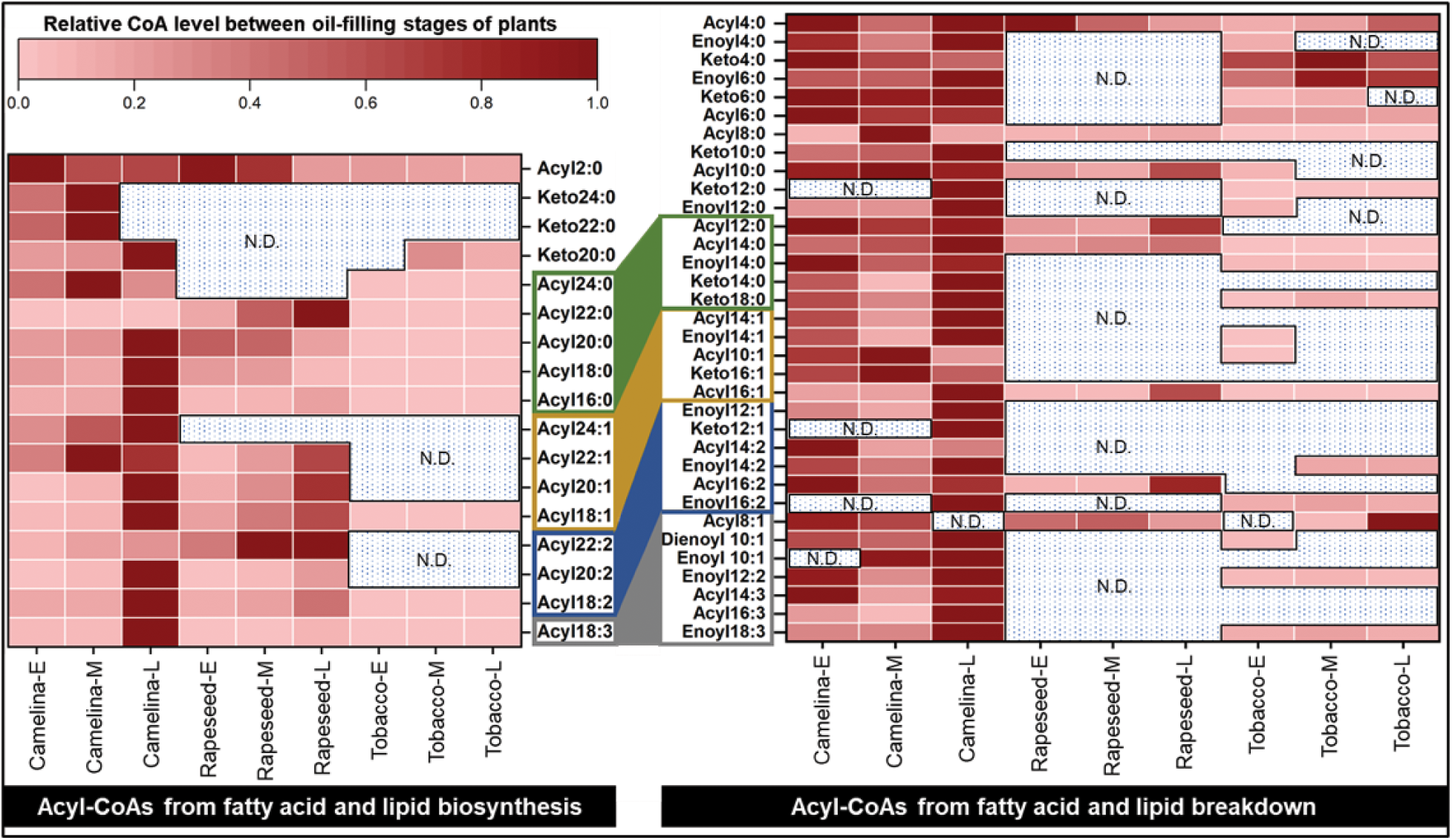
Dynamic fatty acid and lipid metabolism at stages of oil-filing Relative level of fatty acid biosynthesis (left-side) and breakdown-specific (right-side) CoAs from camelina (*Camelina sativa*) and rapeseed (*Brassica napus*) seeds and high-oil tobacco (*Nicotiana tabacum*) leaves in early (E), middle (M), and late (L) development. CoAs representative of a specific unsaturated group of fatty acids are grouped together with same color border. CoAs which overlap in synthesis and breakdown are listed in the synthesis steps. N.D. means ‘not detected’. Heatmaps show mean value (n = 3). t-Tests are presented in Data S2.

The presence of unsaturated short- to medium-chain CoAs in all samples indicated that lipid breakdown during oil-filling is common in oilseeds and engineered leaf tissues. The highest abundance of synthesis- and breakdown-specific CoAs present at later stages of seed development indicated that lipid biosynthesis and breakdown are finely controlled as seed lipid levels approached their maximal value. β-oxidation-specific CoAs were most abundant in camelina seeds. In rapeseed, the detection was limited to a few short-chain acyl-CoAs which were low in abundance, indicating a limited amount of β-oxidation. In contrast to camelina seeds and high-oil tobacco leaves, rapeseed produces less linoleic acid (18:2) and linolenic acid (18:3), and more erucic acid (22:1) (Figure S3) which may reflect a reduced turnover of very long-chain fatty acids. In essence, the presence of fewer long-chain acyl-CoAs for acyl editing to make polyunsaturates in rapeseed could also result in a reduced availability for β-oxidation. When engineered, enzymatic assays and ^14^C feeding experiments in rapeseed inferred an active β-oxidation processes ^17^. Further, a recent ^14^C study in engineered high oil tobacco ^39^ indicated that lipid degradation occurs in tissues actively accumulating lipids; however, the reactions of β-oxidation were not extensively investigated. When we examined concurrent synthesis and breakdown of fatty acids in transgenic high-oil tobacco leaves (Figure 2), the results indicated that triacylglycerol production was unstable ^39,40^. Thus, engineering efforts must consider the collateral effects of breakdown from enhanced lipid production.

### ^18^O-water labeling confirms active β-oxidation processes in oil-filling seeds

As breakdown-specific CoAs were abundant in camelina seeds, this tissue was selected for further studies. Measurements of short- and medium-chain acyl-CoAs provide conclusive evidence that β-oxidation has occurred at some point during development but provides no indication of whether the process is active. To confirm simultaneous lipid synthesis and fatty acid breakdown, the mid-filling stage, when oil accumulation rate is greatest in camelina, was examined through isotopic labeling. ^18^O-labeled water [50% labeled (w/w)] was provided to mid-filling seeds (20 DPA) for 30 min. Rapid incorporation of ^18^O resulted in a mass increase by two (M+2) for 4:0, 8:0, and 14:0 acyl-CoAs, consistent with substitution of ^18^O for naturally abundant ^16^O in acyl-chains during the β-oxidation process (Figure 3A). In 4:0 acyl-CoA, a common breakdown product of all saturated and unsaturated fatty acids (Figure 1B), the presence of 47% ^18^O from 50% isotopic water indicated near complete turnover of the existing short chain acyl-CoA pool within 30 min and signified considerable β-oxidation flux. The higher labeling in 4:0 relative to 8:0 reflected the additional breakdown of tri-unsaturated fatty acids, as 4:0 acyl-CoAs can be produced from all forms of unsaturated fatty acids; but 8:0 can only be produced from mono- and di-unsaturated substrates (Figure 1B). ^18^O signatures in longer-chain acyl-CoAs (≥C16) were likely a consequence of isotope incorporation with thioesterase cleavage of acyl-ACPs during biosynthesis^1,23^.

**Figure 3.**
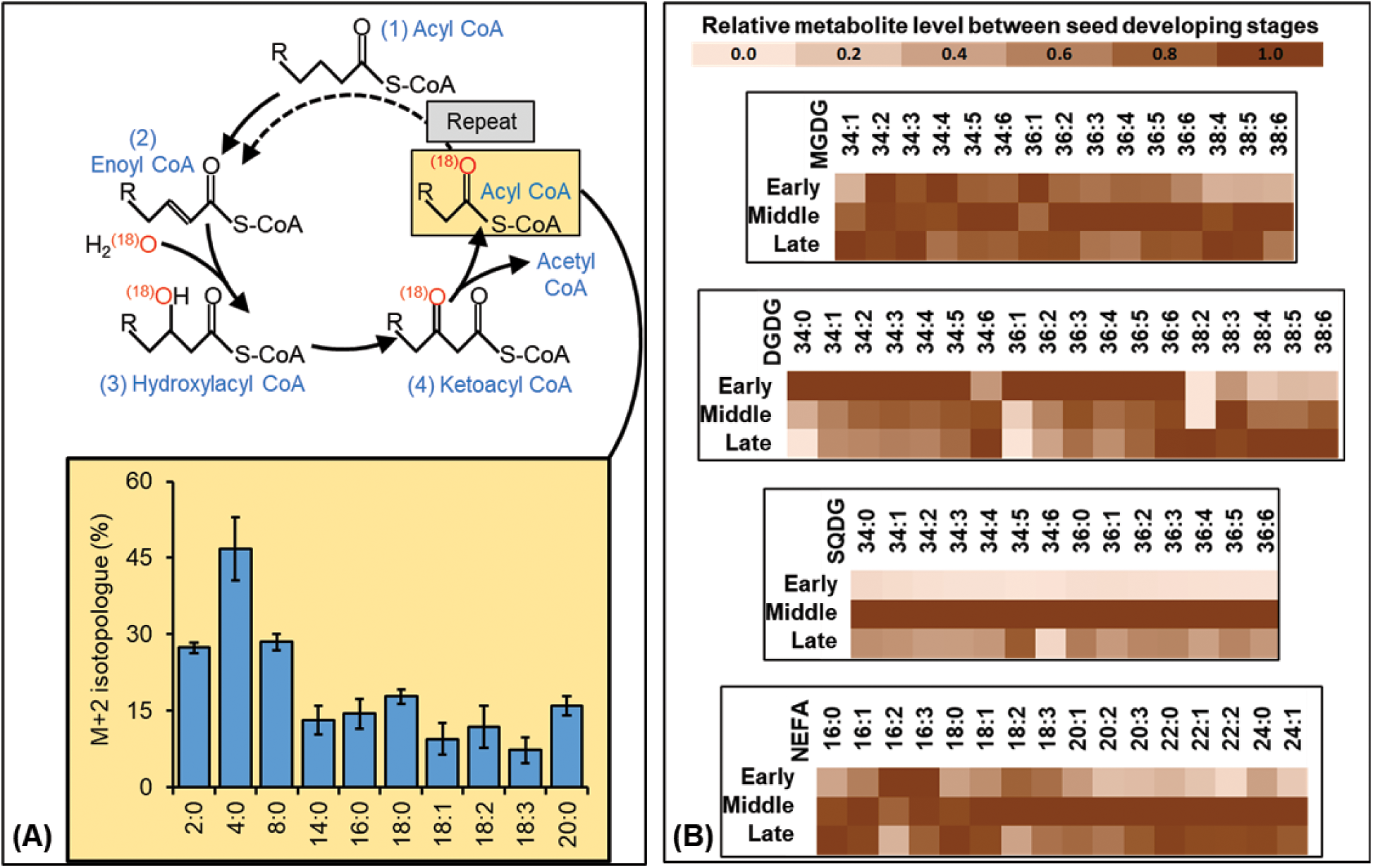
Validation of β-oxidation in oil-filling seeds by labeling study and lipidomics (A) 18O-water labeling of mid-stage camelina seeds for 30 min. Enrichment of M+2 isotopologues are presented (mean ± SD; n=3). (B) Relative level of galactolipids (presented as sum of two fatty acids) and non-esterified fatty acids (NEFA) in developing camelina seeds. Heatmaps show mean values (n = 3). t-Tests are presented in Data S3. Abbreviations (also listed in Data S1): MGDG, monogalactosyldiacylglycerol; DGDG, digalactosyldiacylglycerol; SQDG, sulfoquinovosyl-diacylglycerol; NEFA, non-esterified fatty acid.

### Plastidic galactolipid membranes are turned over as green seeds develop

The breakdown of polyunsaturated fatty acids from lipids was detectable throughout camelina seed development (Figure 2) and supported by changes in the lipid profile (Figure 3B). The presence of 16:1-, 16:2-, and 16:3-containing galactolipids (e.g., 34:4-6 monogalactosyl diacylglycerol or MGDG, digalactosyl diacylglycerol or DGDG, and sulfoquinovosyl diacylglycerol or SQDG) suggested an active prokaryotic lipid biosynthesis pathway operating in seed filling. The abundance of most DGDG and SQDG diminished by late development, consistent with lipid breakdown and possibly chloroplast turnover. Further, the reported biosynthetic rate of combined MGDG and DGDG in mid-filling camelina seeds of 25 pmol h^-1^ embryo^-1^ ^41^ did not result in increasing levels of galactolipids during development (Figure 3B), suggesting that plastid lipid breakdown occurs concomitantly with synthesis. Whereas chlorophyll degradation occurs during maturation and under stress ^42^; the breakdown of DGDG occurred when camelina seeds were green including at early and mid-stages of seed development. The dynamics of breakdown may be associated with maintenance of the photosynthetic complexes including photosystem II or chloroplast thylakoid turnover.

Previously, our group found unconventional acyl-ACPs including 16:1 and 16:3-ACP during seed development in camelina ^24^ which are not part of fatty acid synthesis in plants. Likely, the non-esterified fatty acids (NEFAs) resulting from membrane lipid hydrolysis by lipases are reattached to an ACP backbone ^43^. Here, the results indicated that long-chain NEFAs including 16:1, 16:2 and 16:3 were consistently present throughout seed filling. Fatty acid biosynthesis in chloroplasts and acyl editing in the ER by the activity of phospholipase A2 can produce 16:0 and 18/20/22:x acyl chains; but does not result in polyunsaturated C16 groups ^36^. Conversely, 16:1- to 16:3-containing lipids are produced in chloroplasts by FAD4-8 enzymes and are not thought to undergo acyl editing ^36^. Thus, the presence of 16:1 to 16:3 NEFAs indicated galactolipid degradation and may represent changes in chloroplasts during seed development. Early in development, the abundance of 16:2 and 16:3 NEFAs matched the high proportion of acyl-CoAs derived from di- and tri-unsaturated fatty acid breakdown (Figure 2). These results suggest that galactolipid degradation, followed by fatty acid oxidation, occurs in green oilseeds starting with early stages of seed filling; although the possibility that some fatty acids were used in lipid remodeling cannot be excluded (Figure S4, Table S1). Unlike chloroplast lipids, the degradation of ER-based lipids over development could not be definitively established from lipidomics results (Figure S4). Analysis of available camelina omics information ^44^ indicated a Plastid Lipase 1 (PLIP1) homolog, that is highly expressed in the early-filling stages, may facilitate the degradation of galactolipids (Figure S5, Data S4). Lipases including Oil Body Lipase 1 (OBL1) homologs are highly expressed at early-filling stages supporting that triacylglycerol (TAG) is regularly degraded.

### Peroxisomal oxidation capacity during seed-filling is comparable to that of germination

Fatty acid oxidation in plants has been described in the peroxisomes ^32,45^ where dehydrogenation of acyl-CoA to produce enoyl-CoA in the first step of β-oxidation is catalyzed by acyl-CoA oxidase (ACX) (Figure 1A). Since, both lipidomics and acyl-CoA detection indicated that fatty acid oxidation begins early in the seed-filling, we examined the ACX activity for peroxisomal flux at this stage in comparison to germinating seedlings where oxidation is well-established (Figure 4A). ACX genes utilize distinct substrates of different chain-lengths. ACX1 is specific for long-chain acyl-CoAs (e.g., 16:0), ACX3 utilizes medium-chains (e.g., 8:0) and ACX4 acts on short-chain acyl-CoAs (e.g., 4:0) ^32^. The active breakdown of long to short chain acyl-CoAs was tested with substrates of different chain-lengths. Additionally, 18:3-CoA was used to validate the breakdown of polyunsaturates. The oxidation of long and short chain substrates was comparable in both tissues, indicating the early stages of oil-filling had a similar fatty acid degradation capacity as germinating seedlings. Medium chain oxidation activity was elevated in both tissues but more prominent during germination.

**Figure 4.**
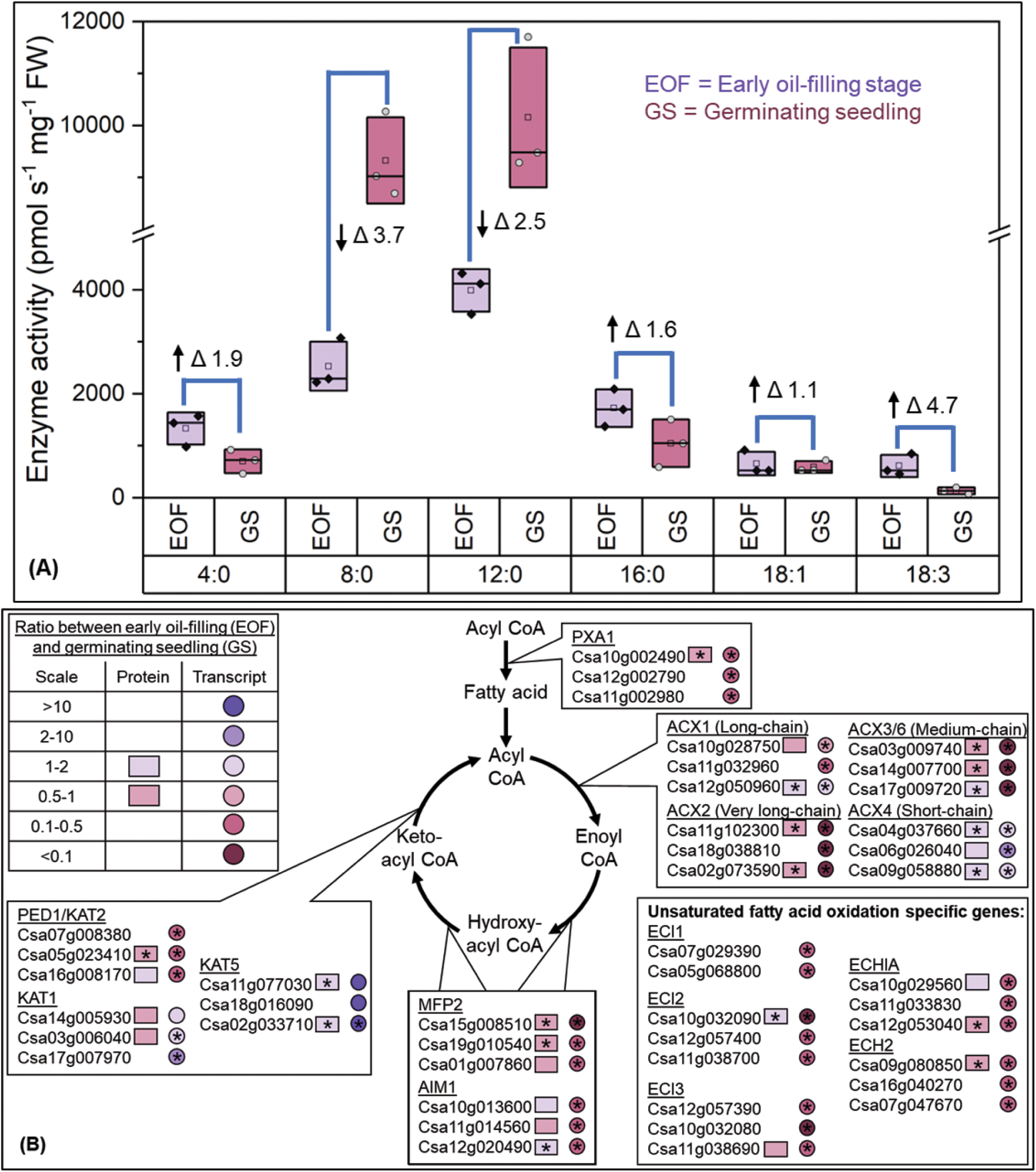
Substantial β-oxidation activity at early oil-filing camelina seeds (A) Six different substrates, namely 4:0, 8:0, 12:0, 16:0, 18:1 and 18:3 acyl-CoA, were provided in independent assays to assess the activity of different acyl CoA oxidases (ACXs) at early oil-filing seeds (EOF) and germinating seedlings (GS) in camelina (n=3). (B) Comparison of protein and transcript abundances of genes related to fatty acid oxidation between EOF and GS. Omics information is presented in Data S5. Statistical significance is indicated by an asterisk (*) at p < 0.05 (n=3).

Potential targets responsible for early seed fatty acid oxidation were assessed through bioinformatic analysis (Figure 4B) of a recently published study ^44^. Among proteins related to β-oxidation, homologs of two keto-acyl CoA thiolases (KAT1 and KAT5) were highly expressed at early-filling stages. Consistent with activity assays, one ACX1 homolog and three ACX4 homologs were higher in abundance and showed higher expression in early-filling seeds. Thus, these homologs of ACX1, ACX4, KAT1 and KAT5 could be potential candidate targets for genetic manipulations to reduce fatty acid breakdown during early oil accumulation in camelina. The reduced expression of medium-chain ACXs at early-filling stages relative to germinating seedling supported the reduced activity for the degradation of 8:0 and 12:0 acyl-CoA substrates. The expression of ACX2, which preferentially utilizes very long-chain substrates (>20C in acyl-chain), was lower supporting our hypothesis that elongated fatty acids are possibly more stable as observed in rapeseed (Figure 2).

### Beta-oxidation occurs during both the day and at night

De novo fatty acid biosynthesis is elevated in light in green tissues including green oilseeds ^46,47^. Alternatively, when leaves are exposed to prolonged darkness, genes associated with lipid degradation are expressed ^48,49^. However, the extent to which fatty acid synthesis and oxidation occurs diurnally, similar to starch degradation in leaves ^50^, is unknown. Developing seeds are thought to stably produce inert storage reserves including storage lipids, thus, the high lipid concentration in seeds may mask the detection of their breakdown during the day. We examined the influence of photoperiod on fatty acid dynamics using a 16-h light, (06:00 to 22:00 h) 8-h dark diurnal cycle. Breakdown-specific acyl-CoA levels were elevated during darkness until morning, reduced at mid-day, and elevated again in the evening in the last phase of the photoperiod (Figure 5A). The results suggest that the diurnal supply of photosynthate impacts lipid degradation, and that fatty acid breakdown may be part of a homeostatic mechanism to ensure sufficient carbon for seed metabolic activities during the night. Additionally, the trend of acyl-CoA consumption and utilization (Figure 5A) suggested that fatty acid biosynthesis (up to 18 carbons) and lipid desaturation were highest early in the day, but fatty acid elongation (20 to 22 carbons) increased later in the photoperiod. Similarly, principal component analysis (PCA) indicated that very long-chain CoAs (C20-24) had distinct metabolism relative to other CoA esters involved in lipid biosynthesis. The analysis of diurnal patterns with PCA indicated that the eigenvectors aligned with expected metabolic processes (Figure 5B), including elevated synthesis (10:00 and 14:00 h) and degradation (02:00 and 06:00 h) that differed from a third period (18:00 and 22:00 h).

**Figure 5.**
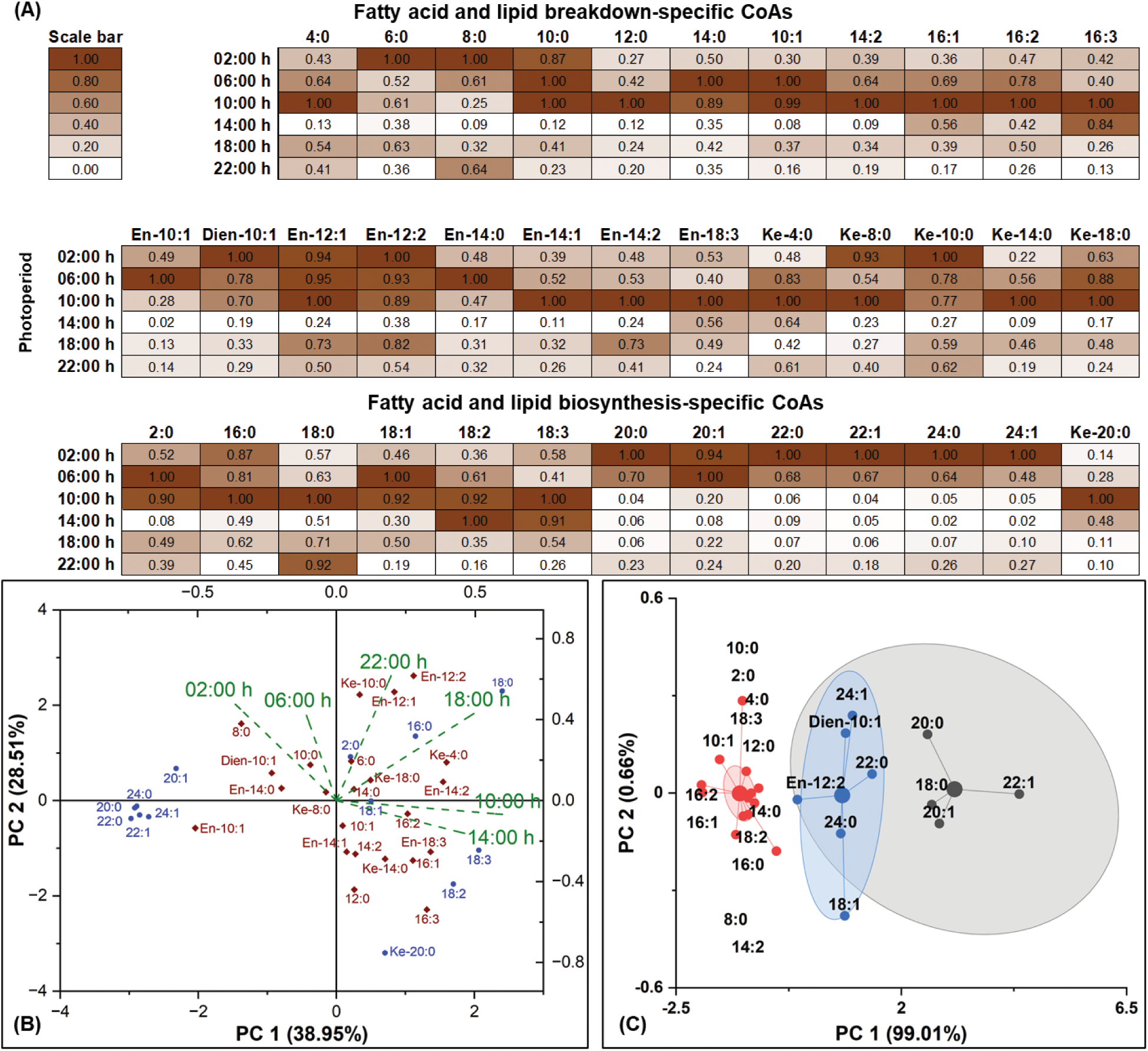
Dynamic fatty acid metabolism over the photoperiod in the middle stage of seed-filling in camelina (A) Heatmap and (B) principal component analysis of relative acyl-CoA abundance in each 4-h window during light (06:00 to 22:00 h) and dark (22:00 to 06:00 h) periods to visualize acyl-CoA changes over 24 h, and clusters of acyl-CoAs and timepoints. Blue, dark red and green colors denote synthesis-specific, breakdown-specific CoAs, and the time of photoperiod, respectively. (C) k-Means clustering analysis of the 13C-touched pool fraction (44) from 8-h of silique culturing in U-13C6 glucose media. n=3 for all three figures. Red, blue, and gray groups represent low, medium, and high 13C-incorporation groups, respectively.

Fatty acid breakdown during the day was further probed with ^13^C glucose labeling using an 8-h silique culturing method ^51^ from 10:00 to 18:00 h. Labeled glucose was metabolized into acetyl-CoA resulting in ^13^C-enriched fatty acids within the 8-h labeling period (Figure 5C). Within the same experimental time, ^13^C was observed in short-chain acyl-CoAs specific for β-oxidation, e.g., 11% ^13^C-containing 4:0 acyl-CoA after correction for natural ^13^C abundance. ^13^C in short-chain acyl-CoAs is only possible when newly produced ^13^C-labeled lipids or fatty acids undergo β-oxidation, unequivocally demonstrating the concomitant biosynthesis and breakdown of fatty acids. The results implied that breakdown was continuous, not unique to nocturnal metabolism. K-means clustering analysis revealed three different clusters that were distinct based on ^13^C description (Figure 5C). The first class consisted of less labeled acyl-CoAs, mostly with specific signatures for β-oxidation (C4-14) and galactolipid breakdown (16:1/2/3), consistent with lipidomics data (Figure 3B). As 4:0-acyl-CoA was at a near isotopic equilibrium by 30-min in the ^18^O-labeling study (Figure 3A), the slower labeling of this pool in the ^13^C study indicated that fatty acid degradation included pre-existing unlabeled lipids in addition to newly synthesized acyl chains. Highly labeled 2^nd^ and 3^rd^ clusters consisted of mostly synthesis-specific CoAs. Compared to 18:1 acyl-CoA, 18:2 and 18:3 belonged to the less-labeled cluster, suggesting that less fatty acid desaturation on phosphatidylcholine (PC) occurs during the experimental period of mid to late photoperiod. Together with this result, a continuous increase of polyunsaturates in abundance in the early light hours, but later decreasing (Figure 5A), suggests that desaturation events occur mostly in the late night to early morning, supporting previous observations in Arabidopsis leaves^52^. Further, the isotope incorporation into 12:2-enoyl- and 10:1-dienoyl-CoA indicates a breakdown of newly formed polyunsaturated fatty acids throughout the day, further confirming the breakdown of lipids as the polyunsaturates are only produced on a lipid backbone (Figure 1).

### Beta-oxidation is a target for engineering

The occurrence of fatty acid breakdown in multiple tissues described here, invited speculation that engineering efforts of the past and present may be thwarted by mechanisms that induce lipid breakdown when plants are altered to produce more oil, perhaps a component of homeostasis. Seeds from high-oil engineered camelina lines were examined to assess whether lipid breakdown altered in response to engineering. Wild-type camelina plant was compared to two transgenic lines with enhanced levels of a α-carboxyltransferase subunit of the acetyl-CoA carboxylase (ACCase) enzyme complex, i.e., full length (FL3-1) and catalytic (CAT A7) lines ^4^, that resulted in higher amounts of seed oil (Figure 6A). Using ^13^C-labeling, isotope incorporation in acyl-ACPs ^24,53^ and -CoAs was measured to compare rates of fatty acid synthesis and breakdown. Both engineered lines produced statistically similar amounts of lipids in their mature seeds (Figure 6A); however, the FL3-1 line showed a higher rate of fatty acid biosynthesis (Figure 6B) and also greater breakdown (Figure 6C). Similarly, a recent study from our group in Arabidopsis showed the increased proteins and transcripts of various peroxisomal oxidation genes when the flux through the de novo fatty acid biosynthetic pathway increased in the double mutant of biotin attachment domain-containing (BADC) proteins, which negatively regulates ACCase ^54^. We postulate that the altered synthesis and breakdown rates could account for an unbalanced production of free fatty acids in the engineered lines that would otherwise be toxic to the cell ^12,17^.

**Figure 6.**
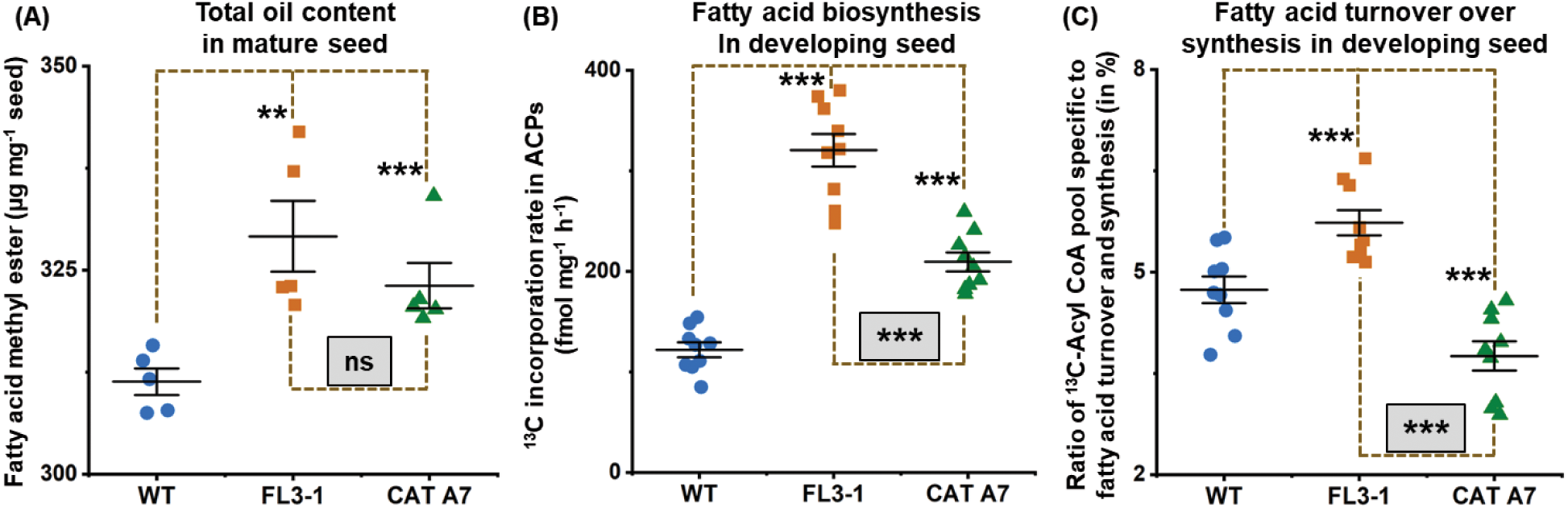
Changes in dynamic fatty acid metabolism in engineered camelina lines (A) Total oil content at maturity based on quantification of fatty acid methyl esters, (B) relative biosynthesis rate at mid-seed filling by measuring 13C ACP pools, and (C) percent breakdown of synthesized fatty acids at mid-seed filling by comparing 13C pools of synthesis-specific long-chain acyl-CoAs (16:0, 18:0 and 18:1) with breakdown-specific short-chain acyl-CoAs (4:0, 8:0 and 10:0) were determined in WT and two transgenic lines (FL3-1 and CAT A7) (Data S6). Data are represented as mean ± S.E.; ** p < 0.03, *** p <0.01, and ns indicates no statistical differences at p=0.05.

Thus, the oil accumulation in transgenics might have been higher if it had not been limited by concomitant breakdown of fatty acids. Enhancing lipid levels with higher synthesis rates (in FL3-1) or lower degradation rates (in CAT A7) suggested that decreasing β-oxidation may help achieve higher lipid levels. We estimated that approximately 4-6% of newly synthesized fatty acids were continuously turned over at the peak of seed filling in WT during the daytime, a conservative estimate that does not account for higher degradation at night (Figure 5A), or enhanced breakdown with early and late filling stages (Figure 2). Lipids are continuously broken down, to repurpose acyl chains as in membrane remodeling that responds to environment or TAG remodeling ^55^. In addition, the current study mechanistically describes β-oxidation of fatty acids, which can have a significant impact on the seed carbon economy and thereby impact development of future oil-rich crops, forcing reconsideration of prevailing assumptions that sink metabolism in seeds is inherently ‘stable’ with implications for breeding and engineering efforts.

## RESOURCE AVAILABILITY

### Lead Contact

Further information and requests for resources and reagents should be directed to and will be fulfilled by the lead contact, Doug K. Allen

### Materials availability

This study did not generate new unique reagents.

### Data and code availability

All data needed to evaluate the conclusions in the paper are present in the paper and/or the Supplementary Materials. Mass spectrometry files, including CoA quantification, labeling measurements in acyl-CoA and acyl-ACP, fatty acyl methyl ester measurements and lipidomics, are deposited in the Zenodo public repository and publicly available upon acceptance of the manuscript for the publication (DOI: 10.5281/zenodo.14393795; 10.5281/zenodo.14396946). This study did not generate any code. Any additional information required to reanalyze the data reported in this paper is available from the lead contact upon request.

## ACKNOWLEDGMENTS

We thank Dr. Tony Larson (University of York) for providing insights on the acyl-CoA method development. We acknowledge the Plant Growth Facility (PGF, Donald Danforth Plant Science Center) for assistance with plant growth and maintenance. We thank Dr. Xue-Rong Zhou from Commonwealth Scientific and Industrial Research Organisation (Australia) for providing the transgenic tobacco lines and Sally Bailey (Donald Danforth Plant Science Center) for technical assistance with measuring acyl-carrier proteins. LC-MS data was acquired in the Bioanalytical Chemistry Facility (BCF) of the Donald Danforth Plant Science Center. The authors greatly appreciate helpful feedback from Drs. Elizabeth (Toby) A. Kellogg, Ian (Max) Moller, Edgar B. Cahoon and Philip D Bates. This work was supported by National Science Foundation grant IOS-1829365, National Science Foundation grant MCB-2242822, Department of Energy Office of Science, Office of Biological and Environmental Research grant DE-SC0023142, Department of Energy Office of Science, Office of Biological and Environmental Research grant DE-SC0022207, U.S. Department of Agriculture-National Institute of Food and Agriculture grant 2021-67013-33778, U.S. Department of Agriculture-National Institute of Food and Agriculture grant 2023-67017-39419, United States Department of Agriculture-Agricultural Research Service (USDA-ARS), and Subterranean Influences on Nitrogen and Carbon (SINC) Center at Donald Danforth Plant Science Center.

## AUTHOR CONTRIBUTIONS

Conceptualization: SK and DKA.; methodology: SK, RBW, and DKA; investigation: SK, PJ, ML, CX, MW, and KLC; visualization: SK and DKA; funding acquisition: AJK, JJT, DX, and DKA; project administration: AJK, JJT, DX, and DKA; supervision: SK and DKA; writing – original draft: SK and DKA; writing – review & editing: SK, PJ, ML, CX, MW, KLC, RBW, AJK, JJT, DX, and DKA

## SUPPLEMENTAL INFORMATION

**Supplementary Document 1:** Text S1: Approximate calculation of annual vegetable oil price globally; Text S2: Beta-oxidation-specific CoAs were sensitively detected in plant samples; Figure S1. Measurement of breakdown-specific CoAs from plant samples in LC-MS/MS; Figure S2. Fatty acyl methyl esters (FAMEs) in different oil-filling stages of camelina and rapeseed seeds and high-oil tobacco leaves; Figure S3. Fatty acids composition (mol %) in late oil-filling stages of camelina and rapeseed seeds and high-oil tobacco leaves; Figure S4. Relative level of membrane and storage lipids in three oil-filling stages of camelina seeds; Figure S5. Relative level of lipase expression at early oil-filling stage over germinating seedling; Table S1. 16:3 fatty acid tail containing membrane and storage lipids present in all three oil-filling stages of camelina.

**Supplementary Document 2:** Data S1: Abbreviations used in the manuscript; Data S2: Statistical analysis of different CoAs between three oil-filling stages of camelina, rapeseed and high-oil tobacco; Data S3: Statistical analysis of different lipid species between three oil-filling stages of camelina; Data S4: Comparison of transcriptomics of different lipase genes in camelina between early-oil filling seeds and germinating seedlings; Data S5: Comparison of proteomics and transcriptomics of fatty acid oxidation related genes in camelina between early-oil filling seeds and germinating seedlings; Data S6: Calculation for fatty acid turnover over synthesis in developing seeds; Data S7: LCMS details for CoA analysis; Data S8: LCMS details for lipidomics analysis.

## STAR★METHODS

### Key resources table

| REAGENT OR RESOURCE | SOURCE | IDENTIFIER |
| --- | --- | --- |
| **Experimental models: Organisms/strains** |  |  |
| *Camelina sativa*: WT | Lab Stock | N/A |
| *Brassica napus*: WT | Lab Stock | N/A |
| Nicotiana tabacum: High-oil line | Commonwealth Scientific and Industrial Research Organisation (Australia) | N/A |
| *Camelina sativa*: High-oil lines | Lab Stock | FL3-1, CAT A7 |
| **Chemicals, peptides, and recombinant proteins** |  |  |
| U-<sup>13</sup>C<sub>6</sub> glucose | Cambridge Isotope Laboratories | CLM-1396 |
| <sup>18</sup>O water | Sigma | 329878 |
| <sup>13</sup>C<sub>3</sub> acyl CoA | Sigma | 655759 |
| <sup>13</sup>C<sub>16</sub> acyl CoA | Sigma | 655716 |
| Methanol | Fisher Scientific | 60-009-55 |
| Methyl tert-butyl ether | Sigma | 34875 |
| UltimateSPLASH ONE mix | Avanti Polar Lipids | 330820 |
| Ammonium acetate | Sigma | 238074 |
| Acetic acid | Sigma | AX0074 |
| Acetonitrile | MedSupply Partners | EM-AX0156 |
| Chloroform | EMD Millipore | CX1058 |
| Sucrose | Research Products International Corporation | S24060 |
| Glutamine | Sigma | 49419 |
| Potassium phosphate dibasic | Sigma | P3786 |
| Potassium Phosphate Monobasic | Sigma | P0662 |
| Isopropanol | Honeywell | 34965 |
| Ammonium bicarbonate | Sigma | A6141 |
| Phosphoric Acid | Sigma | 49685 |
| 2:0 acyl CoA | Avanti Polar Lipids | 870702 |
| 4:0 acyl CoA | Avanti Polar Lipids | 870704 |
| 6:0 acyl CoA | Avanti Polar Lipids | 870706 |
| 8:0 acyl CoA | Avanti Polar Lipids | 870708 |
| 10:0 acyl CoA | Avanti Polar Lipids | 870710 |
| 12:0 acyl CoA | Avanti Polar Lipids | 870712 |
| 14:0 acyl CoA | Avanti Polar Lipids | 870714 |
| 16:0 acyl CoA | Avanti Polar Lipids | 870716 |
| 18:0 acyl CoA | Avanti Polar Lipids | 870718 |
| 16:1 acyl CoA | Avanti Polar Lipids | 870743 |
| 18:1 acyl CoA | Avanti Polar Lipids | 870731 |
| Ammonium sulfate | Sigma | A4915 |
| Flavin adenine dinucleotide (FAD) | Sigma | F8384 |
| Bovine Serum Albumin (BSA) | Sigma | A7030 |
| Triton X-100 | Sigma | X100PC |
| Polyvinylpyrrolidone (PVP) | Sigma | PVP360 |
| Protease inhibitor | Fisher Scientific | 78430 |
| Horseradish peroxidase | Sigma | P8250 |
| p-hydroxybenzoic acid | Sigma | 240141 |
| 4-amino antipyrine | Sigma | 06800 |
| Hydrogen peroxidase | Sigma | 216763 |
| **Deposited data** |  |  |
| Mass Spectrometry Raw Data | Zenodo | 10.5281/zenodo.14393795 |
| Mass Spectrometry Raw Data | Zenodo | 10.5281/zenodo.14396946 |
| **Software and algorithms** |  |  |
| Origin Pro | OriginLab | https://www.originlab.com/ |
| Plant Gramene database | Gramene | http://www.gramene.org |
| The Arabidopsis Information Resource database | Tair | https://www.arabidopsis.org/ |
| **Other** |  |  |
| DB23 column | Agilent Technologies | 122-2332E |
| C8 column | Macherey-Nagel | Custom15M5 |
| Acquity UPLC CSH C18 column | Waters | 186005296 |

## METHOD DETAILS

### Plant growth conditions

All plants were grown at the Donald Danforth Plant Science Center. Greenhouses for camelina and rapeseed were maintained at 22°/20°C (day/night), approximately 50% relative humidity and a 16-h day (06:00 to 22:00 h) and an 8-h night (22:00 to 06:00 h) photoperiod. Natural light varied throughout the seasons; however, sunscreen cloths helped maintain a maximum of 500 µmol m^−2^ s^−1^ on bright summer days with a minimum light level of 250 µmol m^−2^ s^−1^ maintained by a combination of 600-W high-pressure sodium and 400-W metal halide bulbs. Tobacco plants were grown at a 14-h day cycle (08:00 to 22:00 h, 27 °C) and 8-h night cycle (22:00 to 08:00 h, 25 °C). On cloudy days, the light intensity was maintained at a minimum of 500 µmol m^−2^ s^−1^ using supplemental lighting.

### Labeling studies

Isotope tracers were purchased from Sigma-Aldrich (St. Louis, MO, USA). Isotopic glucose (U-^13^C_6_) was fed to siliques containing mid-filling seeds for 4 and 8 h for ACP analysis and for 8 h for acyl-CoA analysis using the established silique culturing technique ^51^. Additionally, intact seeds were supplemented with ^18^O water (H_2_^18^O, 50%) for 30 min in a 50 mM sucrose and glutamine solution based on the imported metabolites in mid-filling camelina seeds ^51^.

### Sample collection

Photoperiod samples were collected every 4 h at 02:00, 06:00, 10:00, 14:00, 18:00, and 22:00 h. All other samples were collected at 10:00 h. For labeling experiments, siliques were harvested around 09.45 h and feeding experiments were initiated around 10:00 h. Samples were flash frozen in liquid nitrogen and stored at −80°C until further analysis.

### Enzyme activity assay

Using a previous method ^32^, protein for ACX activity was extracted from 100 mg (fresh weight) samples of early, mid, and late staged camelina seeds during seed filling, and 2-day old germinating seedlings. ACX enzyme activity was assessed by quantifying the H_2_O_2_ production rate at a wavelength of 500 nm ^32^ in a spectrophotometer (Plate reader Infinite 200 PRO, Tecan, Männedorf, Switzerland). 4:0, 8:0, 16:0, 18:1 and 18:3 acyl-CoAs were used individually as substrates for the enzyme activity assay. To account for potential contamination of CoAs and activity due to existing cellular acyl-CoA pools, blank samples included all components of the reaction mixture except the acyl-CoA substrate. To calculate final enzyme activity rates, the blank value was subtracted from the measured values.

### Extraction of CoAs, ACPs and their measurements in LC-MS

Acyl-, enoyl-, dienoyl- and ketoacyl-CoAs were extracted following previous protocols ^37,56^ with modifications. The first extraction buffer consisted of chloroform and methanol (1:2 v/v) and was stored at −20°C until use. The second extraction buffer was freshly prepared by adding 50 mM KH_2_PO_4_, acetic acid and isopropanol (40:1:20 v/v). A mixture of 1 µM ^13^C_3_ and 0.25 µM ^13^C_16_ acyl-CoAs was used as internal standards in unlabeled samples to account for sample loss for quantification. Approximately 120 mg samples were used for one replicate but were divided into six sub-replicates to increase extraction efficiency. 500 µL of the first buffer and 2 µL of the internal standard mixture were added to each sub-replicate and stirred with beads using a bead beater for 2 min at −20°C. After adding 500 µL of the second buffer, another round of bead beating was performed at 4°C. 10 µL of saturated ammonium sulfate [(NH_4_)_2_SO_4_] was added to the sample and shaken at 4°C for 2 h. After samples were centrifuged at 4°C and 20,000 g for 10 min, the upper phase of all six sub-replicates was pooled in a 15 mL tube for one replicate. Samples were dried in a SpeedVac concentrator and stored at −80°C.

Following a recent protocol ^57^ with modifications, CoAs were measured on a TSQ Altis triple quadrupole mass spectrometer (MS) connected to a Dionex UltiMate 3000 liquid chromatography (LC) system (Thermo Fisher Scientific, Waltham, MA, USA). In the LC, 10 mM ammonium bicarbonate in water was used as mobile phase A and 95% acetonitrile in water as mobile phase B. Analytes were separated using an Acquity UPLC CSH C18 column (1.7 µm × 2.1 mm × 50 mm). Samples were dissolved in 100 µL of solution containing mobile phase A and B and acetic acid (80:15:5 v/v) by sonication for 30 min at 4°C, centrifuged and filtered using 0.8µm PES membrane filters (Sartorius, Göttingen, Germany). Each LCMS run included three injections: one sample injection and two acid wash injections. To reduce carryover in LC column, 25 µL of an acid wash sample consisting of a mixture of phosphoric acid, water, and acetonitrile (5:40:60 v/v) was injected at mobile phase B proportions of 50% and 80% during the column equilibration phase, and the eluent was diverted to waste before entering the MS system. For ionization, the spray voltage was set at 4 kV in positive mode. The ion transfer tube and vaporizer temperature were set at 325°C and 350°C, respectively. The sheath, auxiliary and sweep gas flow rates were set to 45, 7 and 1 arbitrary units, respectively. The LC gradient and compound-dependent MS parameters for CoA are mentioned in Data S7.

Sample extraction and label measurement in acyl-ACPs were performed using previous methods ^24,53,58^. ^13^C-touched pool was calculated by summing all isotopologues (ΣM1-Mn isotopologues, where n is the number of carbon atoms) except the completely unlabeled isotopologue, i.e., M0 ^53^.

### Lipid extraction and measurements in LC-MS

The amount of total lipid was measured by transesterification of fatty acids to fatty acid methyl esters (FAMEs) as previously detailed ^59^. FAMEs were quantified by gas chromatography–flame ionization detection using a DB23 column (30 m, 0.25 mm i.d., 0.25 μm film: J&W Scientific, Folsom, USA) as described ^51^.

Species of lipids were extracted using a variation of the previous protocol ^60^. Homogenized sample material was extracted with a pre-chilled mix of methanol and methyl tert-butyl ether (1:3 v/v) that was spiked with UltimateSPLASH ONE mix (Avanti Polar Lipids, Alabaster, AL, USA) as internal standards. Samples were bead beaten for 30 sec, incubated for 10 min in a shaker at 4°C, and then incubated for another 10 min in an ultra-sonication bath at room temperature. Phase separation was induced by adding 25% methanol in water to each sample tube. After centrifugation, the upper organic layer was aliquoted to a new tube and dried down in a SpeedVac concentrator. Samples were analyzed in LC-MS using a Q Exactive mass spectrometer coupled to a Dionex UltiMate 3000 UHPLC system (Thermo Fisher Scientific). A C8 column (5 µm x 150 mm x 0.5 mm) was used for chromatographic separations. Mobile phase A consisted of water with 1% 1 M ammonium acetate and 0.1% acetic acid, and mobile phase B consisted of 7:3 acetonitrile: isopropanol with 1% 1 M ammonium acetate and 0.1% acetic acid. The mass spectrometer acquired both positive and negative mode full scan mass spectra (MS1) from a single injection using polarity switching. For ionization, the spray voltage was set at 4 kV for positive mode and 3.9 kV for negative mode. The capillary and auxiliary gas heater temperature were set at 250°C and 60°C, respectively. The declustering potential (RF lens) was set to 50 V. The sheath and auxiliary gas flow rates were set to 15 and 5 arbitrary units, respectively. LC gradient data dependent MS1 and MS2 parameters and quantitative method details of measured lipid species are mentioned in Data S8.

### Bioinformatics

Proteins specific to lipase activity and straight-chain fatty acid breakdown were selected from Arabidopsis. Homologous proteins in camelina were identified using the Gramene database ^61^. The extracted homologous proteins were annotated with gene symbols and descriptions based on the TAIR11 protein database ^62^ to ensure consistent and accurate functional characterization. The abundance and expression of identified camelina protein candidates were extracted from a previous study and reanalyzed for comparison between different developmental stages ^44^.

