## Supplementary Document for "Persistent fatty acid catabolism during plant oil synthesis"

**The PDF file includes:**

Supplementary Text S1 to S2

Figures S1 to S5

Table S1

References used in this Supplementary Document

### SUPPLEMENTARY TEXT:

#### Text S1. Approximate calculation of annual vegetable oil price globally

- The global production of major vegetable oil was ~218 million tons in 2022/23 (data collected from Oilseeds: World Markets and Trade, March 2024, United States Department of Agriculture Foreign Agricultural Service).
- The global vegetable oil price was ~$1650/ton in 2022/23 (price value calculated based on price index information from Oilseeds, Oils and Meals. Monthly price update no. 176, March 2024, Food and Agriculture Organization of the United Nations).
- The total global oil price was ~$400 billion in 2022/23.
- 1% of global oil production is equivalent to ~$4 billion per year.

#### Text S2. Beta-oxidation-specific CoAs were sensitively detected in plant samples

To sensitively quantify different key intermediates of the β-oxidation process, the previous extraction protocol and LC-MS / MS quantification method were optimized using developing seed and leaf samples. Following the previous extraction method [1], reduced loss of acyl-CoAs during extraction was achieved using approximately 25 mg of biomass in each replicate as suggested earlier [2], however shorter chain acyl-CoAs except acetyl-CoA were confined to background MS noise. To detect low-abundance metabolites without compromising extraction efficiency, six sub-replicates containing 15-25 mg biomass each were extracted separately and combined before vacuum drying to form a biological replicate. A few steps of the original protocol (e.g., use of bovine serum albumin and saturated petroleum ether) were omitted because they were either not essential for LC-MS based analysis or to avoid sample loss.

The carryover of long-chain acyl-CoAs was a main problem in LC-MS measurement and can be solved by using multiple autosampler-injections of phosphoric acid at the end of each run [3]. This acid has an adverse impact on the negative ionization mode of MS; therefore, the number of acid wash injections was optimized and reduced from ten injections of 50 µL each to two injections of 25 µL each. Acids were injected at 50% and 80% buffer B conditions during the 16 min of cleaning and equilibration time, resulting in the absence of carryover of acyl-CoAs and phosphate ions. Further, the pool size of acyl-CoAs in plant samples is typically low (fmol/mg range [1]); hence, the initial LC gradient to capture CoAs was altered to reduce ion competition during electrospray ionization and to increase MS sensitivities. The compounds were eluted slowly and separated during the 12 min of chromatographic detection (Figure S1A). Acyl-CoAs having a close molecular weight (e.g., 10:0 vs 10:1) were distinctly measured to avoid the interference by isotopologues in quantitative and labeling analysis; for instance, the separation of 10:0 and 10:1 is necessary since the M2 isotopologue of 10:1 and M0 isotopologue of 10:0 have the same molecular weight (Figure S1B). When a standard was not available, an acyl-CoA was screened based on the unique neutral loss of 507 Da during fragmentation and retention time trend, e.g., 16:2 was predicted to be detected before 16:1 as similar to the detection of 18:2 and 18:1 standard.

Using the optimized methods, we measured breakdown-signature CoAs in germinating camelina seedlings (Figure S1C). As acetyl-CoA (C2:0) is the end product of β-oxidation, this metabolite had the highest abundance in germinating seedlings. Butanoyl-CoA (C4:0) is the only common saturated acyl-CoA which is produced in the breakdown of saturated, mono- and poly-unsaturated fatty acids (Figure 1B) and had higher abundance in germinating seedlings compared to other saturated acyl-CoAs containing higher carbon-chain lengths (Figure S1C). Further, the presence of unsaturated acyl-, enoyl-, and dienoyl-CoAs, which were not previously measured in seedlings, confirmed the breakdown of lipogenic polyunsaturated fatty acids during germination.

**Supplementary Figures:**

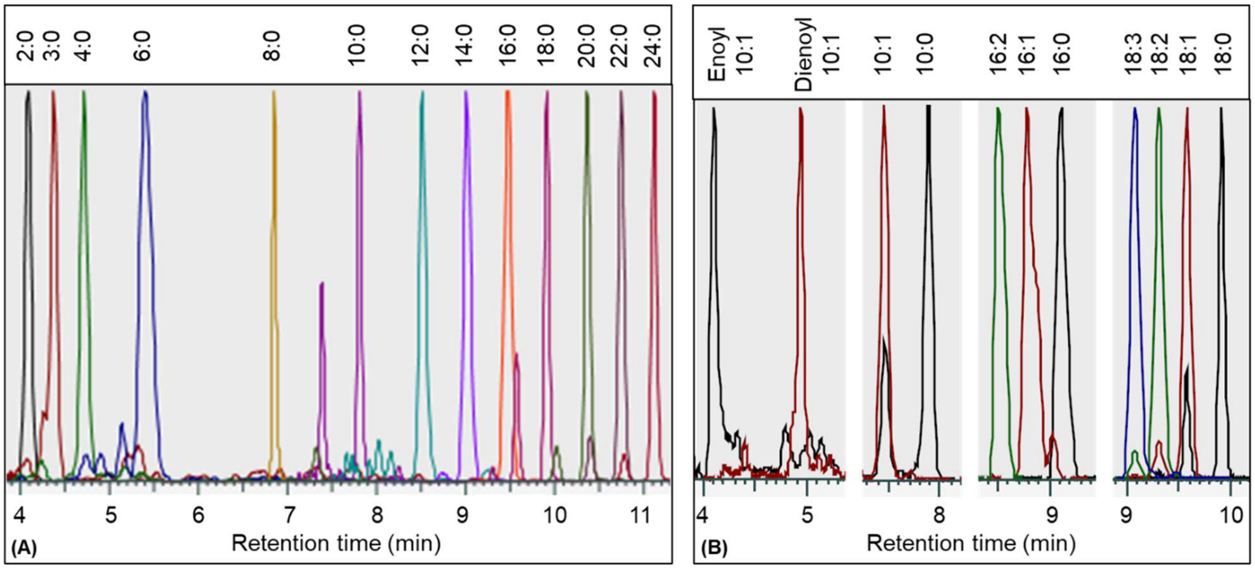


| (C) β-oxidation specific CoAs | Metabolite abundance (fmol/mg) |
|---|---|
| **Acyl 2:0** | 126.3 ± 37.6 |
| **Acyl 4:0** | 10.6 ± 2.4 |
| **Acyl 6:0** | 0.5 ± 0.1 |
| **Acyl 8:0** | 0.8 ± 0.1 |
| **Acyl 10:0** | 1.4 ± 0.4 |
| ***Acyl 10:1** | 2.6 ± 1.1 |
| **Acyl 12:0** | 0.6 ± 0.4 |
| **Acyl 14:0** | 3.1 ± 1.1 |
| ***Acyl 14:2** | 3.2 ± 1.5 |
| ***Dienoyl 10:1** | 1.7 ± 0.4 |
| ***Enoyl 10:1** | 0.4 ± 0.4 |
| ***Enoyl 12:1** | 3.9 ± 0.7 |
| ***Enoyl 12:2** | 2.1 ± 0.7 |
| ***Enoyl 14:0** | 2.3 ± 1.5 |
| ***Enoyl 14:1** | 9.6 ± 6.3 |
| ***Enoyl 14:2** | 2.8 ± 0.1 |
| ***Keto 4:0** | 10.7 ± 2.9 |
| ***Keto 8:0** | 0.5 ± 0.01 |
| ***Keto 10:0** | 1.7 ± 0.8 |
| ***Keto 14:0** | 2.7 ± 0.7 |

**Figure S1. Measurement of breakdown-specific CoAs from plant samples in LC-MS/MS.** (A) Chromatogram of saturated CoAs measured from extracted camelina seed sample. (B) Separation of the unsaturated forms from the saturated form of acyl-CoAs with the same number of carbon atoms in the acyl chain. (C) Quantification of breakdown-specific CoAs in 2-days old germinating camelina seedlings. Quantification was performed based on the standard curve, except for CoAs marked with an asterisk (*) for which no standards were available. In these cases, the quantification of unsaturated CoAs was performed based on the standard curve of 16:1 acyl-CoA, and the quantification of the keto and enoyl forms of saturated CoAs was performed based on their acyl form.

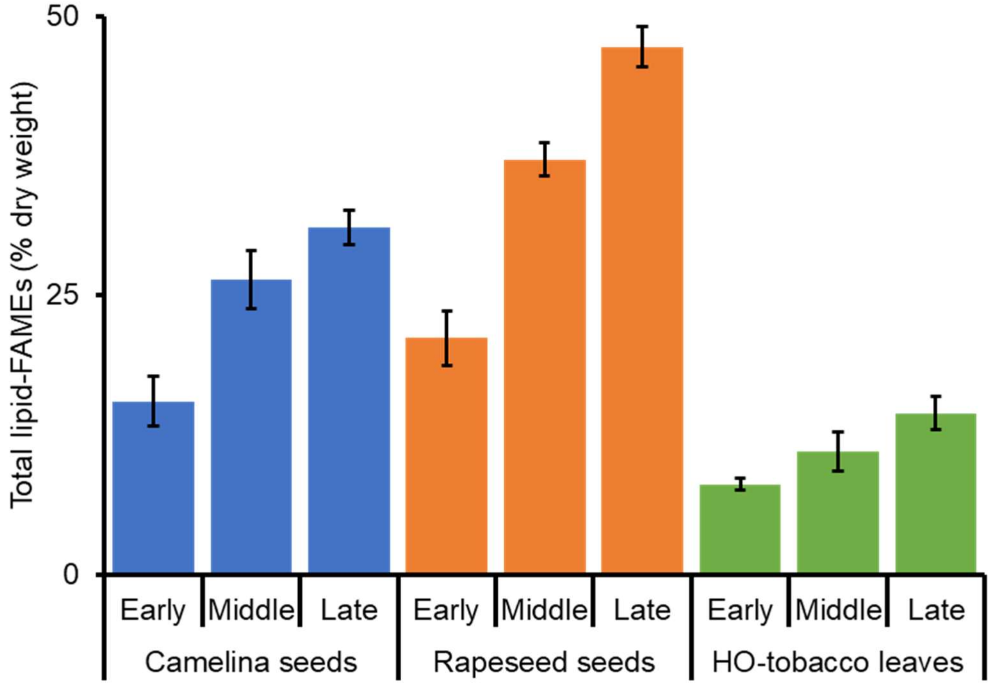


**Figure S2. Fatty acyl methyl esters (FAMEs) in different oil-filling stages of camelina and rapeseed seeds and high-oil tobacco leaves.** (mean ± SD; n=3)

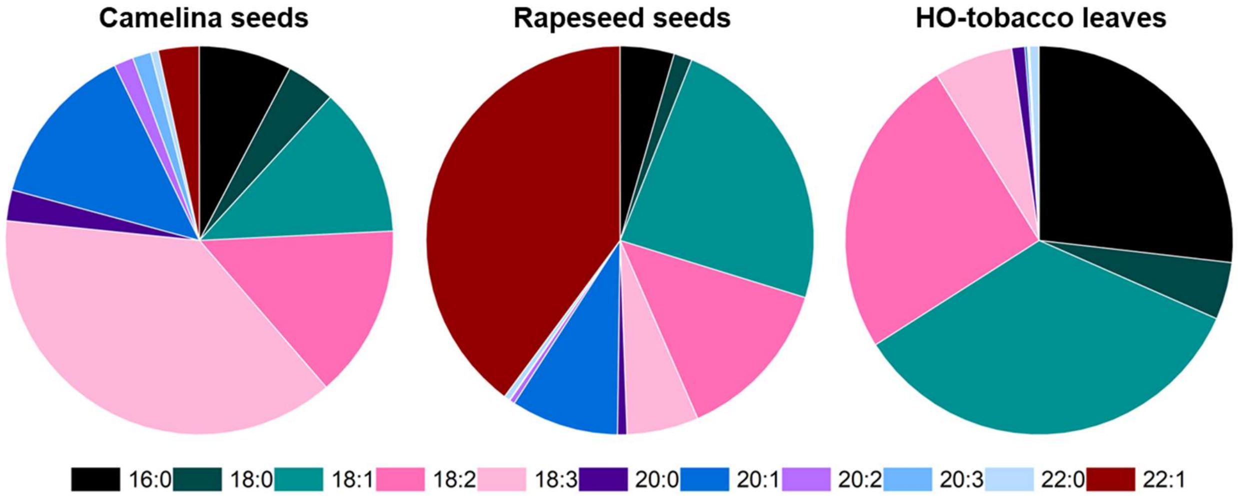


**Figure S3. Fatty acids composition (mol %) in late oil-filling stages of camelina and rapeseed seeds and high-oil tobacco leaves.** (mean; n=3)

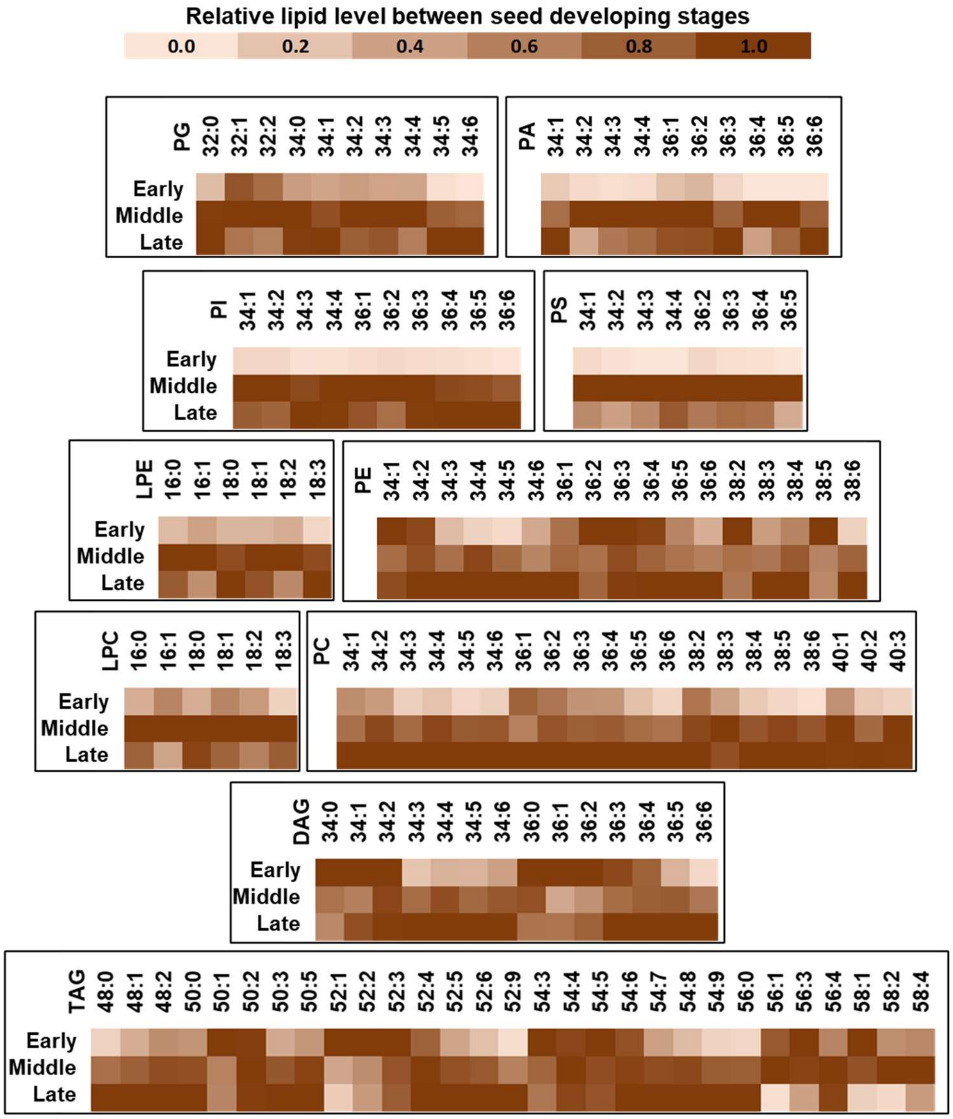


**Figure S4. Relative level of membrane and storage lipids in three oil-filling stages of camelina seeds.** Heatmaps include mean value (n = 3). T-tests are presented in Data S3. There is no evidence from the lipidomics that the 16:2 and 16:3 end up on ER lysolipids (lyso-phosphatidylcholine or LPC and lyso-phosphatidylethanolamine or LPE), although other ER lipids containing 16:3 were identified (Table S1), suggesting some extent of 16:3 based lipid remodeling in addition to β-oxidation. Alternatively, the presence of 16:1 in ER-lysolipids could indicate the production of 16:1 ACP by a non-specific stearoyl-ACP desaturase [4]. No NEFAs for chain lengths shorter than 16-carbon were detected (Figure 2C), further supporting the involvement of C4-14 chain lengths in β-oxidation. Additionally, some membrane lipids such as phosphatidylglycerol (PG), phasphatidic acid (PA), phosphatidylserine (PS), LPC, and LPE were increased until the mid-filling stage but decreased in the late-filling stage. This suggests possible late-stage membrane lipid breakdown but could also be due to lipid remodeling. Phosphatidylinositol (PI), PE and PC were increased at later stages. Similarly, most di- and tri-acylglycerols (DAG and TAG) were elevated during development.

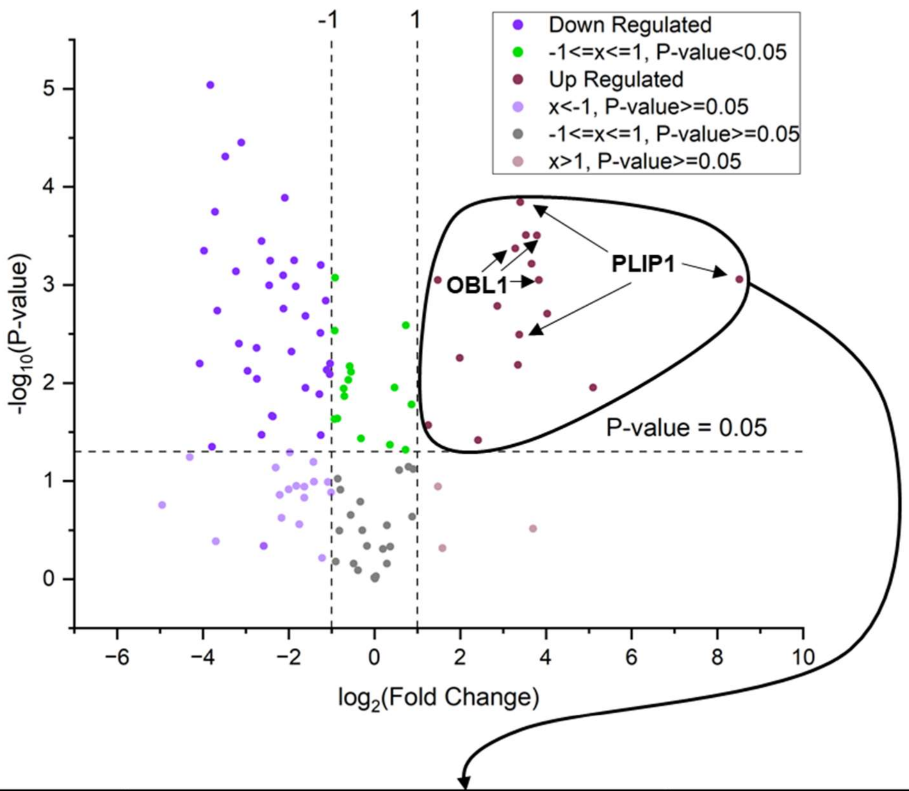


| Gene ID | Gene Description | Specific/Preferred Substrates |
|---|---|---|
| Csa01g038410 | PHOSPHOLIPASE A2-ALPHA (PLPA2-ALPHA) | Linoleoyl acyl chain at sn-2 position of phosphatidylcholine (e.g., PC -> LPC) |
| Csa15g059780 | | |
| Csa19g040900 | | |
| Csa07g060440 | NON-SPECIFIC PHOSPHOLIPASE C2 (NPC2) | Phosphatidylcholine (e.g. PC -> DAG) |
| Csa01g016910 | OIL BODY LIPASE 1 (OBL1) | Triacylglycerol |
| Csa15g018720 | | |
| Csa19g021040 | | |
| Csa05g093810 | PLASTID LIPASE1 (PLIP1) | Galactolipids (e.g., MGDG -> LMGDG) |
| Csa07g002960 | | |
| Csa16g003810 | | |
| Csa10g007060 | PHOSPHOLIPASE-LIKE PROTEIN 4 (PEARLI 4) | - |
| Csa11g004610 | PHOSPHOLIPASE A IVC (PLAIVC) | Linoleoyl acyl chain at sn-2 position of phosphatidylcholine (e.g., PC -> LPC) |
| Csa12g004350 | | |
| Csa02g064320 | PHOSPHOLIPASE C1 (PLC1) | Phosphatidyl-myo-inositol bisphosphate |
| Csa18g031850 | | |
| Csa18g035700 | PROTEINS PHOSPHOLIPASE 3A (PGAP3A) | Glycosylphosphatidylinositol |

**Figure S5. Relative level of lipase expression at early oil-filling stage over germinating seedling.** Details are in Data S4.

**Supplementary Table:**

**Table S1. 16:3 fatty acid tail containing membrane and storage lipids present in all three oil-filling stages of camelina**

| Lipid species | m/z | Adducts | Identified tails |
|---|---|---|---|
| PC 34:6 | 786.5 | [M+OAc]- | 16:3 and 18:3 |
| DAG 34:4 | 606.5 | [M+NH4]+ | 16:3 and 18:1 |
| DAG 34:5 | 604.5 | [M+NH4]+ | 16:3 and 18:2 |
| DAG 34:6 | 602.5 | [M+NH4]+ | 16:3 and 18:3 |
| TAG 50:5 | 842.7 | [M+NH4]+ | 16:3, 16:0 and 18:2 |
| TAG 52:9 | 862.7 | [M+NH4]+ | 16:3, 18:3 and 18:3 |

**REFERENCES (used in this supplementary document)**


1. Larson, T.R., and Graham, I.A. (2001). Technical Advance: a novel technique for the sensitive quantification of acyl CoA esters from plant tissues. Plant J. Cell Mol. Biol. *25*, 115–125. https://doi.org/10.1046/j.1365-313x.2001.00929.x.

2. Haslam, R.P., and Larson, T.R. (2021). Techniques for the Measurement of Molecular Species of Acyl-CoAAcyl-CoA in Plants and MicroalgaeMicroalgae. In Plant Lipids: Methods and Protocols Methods in Molecular Biology., D. Bartels and P. Dörmann, eds. (Springer US), pp. 203–218. https://doi.org/10.1007/978-1-0716-1362-7_12.

3. Pearce, R.W., Kodger, J.V., and Sandlers, Y.I. (2022). A liquid chromatography tandem mass spectrometry method for a semiquantitative screening of cellular acyl-CoA. Anal. Biochem. *640*, 114430. https://doi.org/10.1016/j.ab.2021.114430.

4. Drissner, D., Kunze, G., Callewaert, N., Gehrig, P., Tamasloukht, M., Boller, T., Felix, G., Amrhein, N., and Bucher, M. (2007). Lyso-phosphatidylcholine is a signal in the arbuscular mycorrhizal symbiosis. Science *318*, 265–268. https://doi.org/10.1126/science.1146487.
